# Artificial light at night enhances the defensive behavior in aquatic shredders under predation risk

**DOI:** 10.1101/2024.10.16.618652

**Authors:** Zahira Rufaida, Magdalena Czarnecka

## Abstract

Caddisfly larvae (Trichoptera) are mostly nocturnal aquatic invertebrates that contribute to the decomposition of coarse organic matter, such as plant litter in streams. Artificial light at night (ALAN) can interfere with their behavior, particularly in the presence of predation threat. We investigated the effects of various types of ALAN on *Lepidostoma hirtum*, a common caddisfly species in running waters. We tested its behavior (activity and tendency to shelter) in darkness, as well as under white warm LED light and HPS (sodium lamp) light (2 lux), with and without the presence of predation cues (crushed conspecifics). We found that in darkness caddisflies did not change their behavior in the presence of predation threat. When exposed to ALAN only, caddisflies decreased their activity compared to dark nights, regardless of the type of light. The presence of both predation threat and ALAN enhanced a defensive behavior in caddisflies, especially under LED light. They not only reduced their activity, but also spent more time in shelter, compared to other treatments. This study highlights that impact of ALAN on the behavior of nocturnal aquatic invertebrates may vary depending on the spectral composition of light and may be intensified in the presence of predation threat.

## Introduction

Artificial light at night (ALAN) has been affecting the natural environment for many years and its effect is constantly increasing due to progressive urbanization (Kyba et al. 2011; Papalambrou and Doulos 2019; Kostrakiewicz - Gieralt et al. 2022). Falchi et al. (2016) revealed that around 83% of the world’s population and over 99% in the United States and Europe are exposed to light-polluted skies, with a rapid growth observed, particularly in Asian countries characterized by extreme night brightness and poor night sky visibility. Sources of disturbance include direct light emission from street lighting, households and industry, as well as skyglow resulting from artificial light scattering and reflecting back to earth through the atmosphere (Gaston et al. 2013; Kyba et al. 2011; Davies et al. 2013).

One of the main effects of ALAN on organisms is the disruption of natural light cycles (Longcore and Rich 2004), which may significantly affect their physiology (Davies et al. 2014; Kupprat et al. 2020b) and behavior (Czarnecka et al. 2019; 2022; Duarte et al. 2019). ALAN may influence animal behavior at night in various ways. Many animals rely on light for navigation, and ALAN can alter their movement patterns either by attracting them directly to the light or by masking natural light sources used for orientation due to skyglow (Gaston et al. 2013). Effects of ALAN can differ between diurnal and nocturnal species. Diurnal and crepuscular animals tend to become more active at night in the presence of ALAN, exploiting the newfound night light niche (de Jong et al. 2016; Russ et al. 2015; Dwyer et al. 2013), while nocturnal animals often become less active at night (Lewanzik and Voigt 2014; Rotics et al. 2011). This pattern has been observed in various animals, including insects (van Langevelde et al. 2011), birds (Gauthreaux and Belser 2006), amphibians (Buchanan 2006; van Grunsven 2017), reptiles (Witherington and Bjorndal 1991). However, the majority of studies on ALAN have investigated its effects on terrestrial organisms, leaving a gap in our understanding of how ALAN affects behavior of aquatic species. Moreover, we know relatively little on the impact of the quality of artificial light on animal behavior. Spectral composition of ALAN differs from natural light sources, such as moonlight (Kelber et al. 2017), and this can also be a source of disturbance. Previous studies have shown that LED light containing a significant proportion of blue light can cause stress and promote light avoidance behavior in aquatic invertebrates (Manríquez et al. 2019; Fischer et al. 2020), while high-pressure sodium (HPS) lamps emitting yellow-orange light are considered less harmful for wildlife (Pawson and Bader 2014; Davies et al. 2017; Davies and Smyth 2018).

The animal response to ALAN may also be modified when predators appear. It is well known that the physical presence of a predator or traces of its kairomones in the environment may trigger defensive behavior in a potential prey. Aquatic species employ various strategies when faced with predation pressure, such as escaping, concealment, dispersing or forming larger groups to minimize the risk of capture (Chivers and Smith 1998; Brown 2003; Ferrari et al. 2010). Since illuminated areas may be perceived as potentially dangerous, especially by nocturnal species, it is possible that defensive behaviors may increase in the presence of predation cues. As a result, ALAN can strongly determine the outcome of predator-prey interactions, as well as influence interactions between conspecifics (Miller et al. 2017).

In this study, we investigated behavioral responses of *Lepidostoma hirtum* (Trichoptera), a common caddisfly species in lowland streams to low-intensity ALAN (2 lx) differing in the spectral composition (high pressure sodium HPS light and white warm LED light). We tested the behavior of caddisflies both in the presence or absence of predation cues, provided by injured conspecifics.

*L. hirtum* is a nocturnal species and occurs in relatively shallow habitats which may be particularly susceptible to artificial light exposure throughout the year (Brüning et al. 2018). *L. hirtum* plays a significant role in the cycling of organic matter and energy flow (Cummins 1973), as it exhibits a high capability to fragment coarse particulate organic matter (leaf litter) in streams and thus accelerates decomposition (Azevedo-Pereira et al. 2006). Individuals of *L. hirtum* often occur in groups, colonizing leaf litter, woody debris or stones in slow flowing streams (Schmidt-Kloiber and Hering 2015; Graf et al. 2023). They also construct portable cases made of sand grains that provide physical protection against predator attack (Hansell 1972; Ferry et al. 2013).

In this experiment, we observed the behavior of caddisflies under different light conditions (darkness, HPS and LED light) and estimated time spent in movement and shelter use, as well as their tendency to form groups, both in the presence or absence of predation threat. We assumed that exposure to ALAN would modify behavior of *L. hirtum* and reduce its activity, because this nocturnal species may associate increased light levels with higher predation risk. This effect would be particularly pronounced in the presence of predation cues. We also expected that caddisflies would be more sensitive to LED light containing a high proportion of blue light than to the yellow-orange light produced by HPS lamps.

## Methods

### Animal collection and housing

In March 2022, we collected caddisflies *Lepidostoma hirtum* from Zielona Struga stream (52.997208 N, 18.450469 E), situated in a forest approximately 11 km from Torun, a city with around 200,000 inhabitants. The location was chosen due to minimal exposure to light pollution. Using a hand net, we collected the specimens from the streambed, together with fine woody debris and decaying leaves. The caddisflies were transported to the laboratory and housed in an 80 L aquarium in an air-conditioned room. The aquarium contained settled tap water, and its bottom was covered with fine wood and leaves obtained from the stream, providing food for caddisflies. The animals were maintained in aerated water at a constant temperature of 15°C and exposed to natural photoperiod through windows. The caddisflies were acclimated to laboratory conditions for a week before the start of the experiment.

### Experimental setup

We investigated the impact of ALAN on the behavior of caddisflies, following a modified protocol from Czarnecka et al. (2021). The experiment was conducted in a darkened room. We mimicked daylight conditions with incandescent lighting provided from 07:00 to 19:00. To simulate light-polluted conditions, we installed HPS (50W, 4500 lm, 2000 K, SON-T Philips) and LED (5.5W, 470 lm, 2700 K, Philips) lamps above the experimental arenas. Light intensity was adjusted to achieve 2 lux at the water surface. This light level corresponds to the values observed in waters in urban areas (Perkin et al., 2014). Control treatment was performed in darkness (<0.01 lx). Each experimental arena consisted of a round white dish with a diameter of 10 cm. Each dish had its own aeration system and was placed in a shallow tank filled with water, which was continuously cooled by aquarium coolers (Teco R20; Teco S.l.r. Ravenna, Italy). This cooling system maintained the water temperature at 15°C throughout the experiment. We also placed three stones in each dish (in a random position) to provide shelter for the caddisflies. Digital cameras were installed above the experimental arenas to record the caddisfly behavior. Observations in darkness were conducted using a low-power infrared lamp.

### Experimental procedure

We introduced four caddisflies into each dish before the experiment started. The caddisflies were acclimated to the conditions in the experimental arena for 15 minutes. We then started recording their behavior using cameras. Behavioral observations lasted 45 minutes.

To assess the interaction between ALAN exposure and potential predation risk, just before the experiment we prepared a mixture which served as a source of cues indicating the presence of predator. We crushed two new caddisflies using mortar and pestle. Then we added 10 mL of distilled water, homogenized the mixture and filtered it through a fine mesh to remove caddisfly residue. One portion of the solution (10 mL) was supplied to each dish before we introduced living caddisflies and started the experiment.

After completing the trials, we analysed the caddisfly behavior from video recordings using Ethovision 10.1 software (Noldus Information Technology bv, Wageningen, The Netherlands). We estimated the following variables based on the ethogram: 1) the percentage of time spent in movement, 2) total distance covered by Trichoptera (cm), 3) the percentage of time spent in shelter, 4) the mean and minimum distances between caddisflies (to estimate their tendency to group formation). In total, we completed 96 trials consisting of three light treatments, with and without the predator cues. We conducted 16 replicates in each treatment.

### Data analysis

We conducted data analysis using PS Imago Pro 9.0 (IBM SPSS Statistics 29). Since data were mostly normally distributed and variances within groups were homogeneous, we used General Linear Model (GLM) to analyse the effect of different light conditions (darkness, HPS or LED light) and presence or absence of predation cues on time spent in movement and distance covered by caddisflies, time spent in shelter, and the mean and minimum distances between caddisflies (indicating grouping behavior). Significant effects from the models were further analysed using sequential Bonferroni corrected post-hoc tests.

## Results

We found that both exposure to ALAN and predation cues significantly affected the behavior of *L. hirtum*. Generally, caddisflies exhibited low activity, however we found some differences in time spent moving between light treatments. In the absence of predation cues, *L. hirtum* reduced its activity in LED and HPS light compared to darkness (Table 1A, Table 2, Fig. 1A), but did not increase time spent in shelter (Table 1B, Table 2, Fig. 1B). We found no impact of predation threat on the activity of caddisflies and shelter use in darkness (Fig. 1A, Fig. 1B). Caddisflies exposed to ALAN under predation risk decreased their activity compared to darkness (Fig. 1A), and they maintained it at a similar level as in the ALAN treatment without predation cues. However, we observed differences in the shelter use when *L. hirtum* was exposed to predation cues, depending on the type of light. Caddisflies in LED light spent more time in refuge than in HPS light and darkness (Table 1B, Table 2, Fig. 1B). They also spent less time in shelter in HPS light when exposed to predation risk than in the absence of predation cues (Fig. 1B). Neither predation cues nor light conditions had any effect on the distance moved by caddisflies (Table 1C), as well as on their grouping behavior (mean distance and minimum distance between individuals) (Table 1D,E).

**Table 1.** Effects of light type (darkness, HPS, LED) and predation cues on time spent in shelter (A), total distance (cm) covered by caddisflies (B), mean (C), and minimum distance (cm) (D) between individuals of caddisflies (General Linear Model). Statistically significant results are indicated by asterisks.

| Dependent variables | Effects | df | MS | F | P |
| --- | --- | --- | --- | --- | --- |
| A. Time spent in movement (%) | Light | 2 | 6.428 | 13.982 | <0.001* |
|  | Signal | 1 | 1.201 | 2.613 | 0.11 |
|  | Signal*Light | 2 | 0.001 | 0.003 | 0.997 |
| B. Time spent in shelter (%) | Light | 2 | 2495.549 | 8.895 | <0.001* |
|  | Signal | 1 | 668.629 | 2.383 | 0.126 |
|  | Signal*Light | 2 | 1311.268 | 4.674 | 0.012* |
| C. Mean distance covered by caddisflies (cm) | Light | 2 | 195411.976 | 2.592 | 0.08 |
|  | Signal | 1 | 244176.491 | 3.238 | 0.075 |
|  | Signal*Light | 2 | 65596.395 | 0.87 | 0.422 |
| D. Mean distance between individuals (cm) | Light | 2 | 0.146 | 0.256 | 0.775 |
|  | Signal | 1 | 0.976 | 1.712 | 0.194 |
|  | Signal*Light | 2 | 0.28 | 0.492 | 0.613 |
| E. Minimum distance between individuals (cm) | Light | 2 | 0.044 | 0.581 | 0.562 |
|  | Signal | 1 | 0.197 | 2.597 | 0.111 |
|  | Signal*Light | 2 | 0.01 | 0.136 | 0.873 |

**Table 2.** Post-hoc comparisons for significant effects of the light type (darkness, HPS, LED) and predation cues on time spent in movement and time spent in shelter by caddisflies. The main models are shown in Table 1. Statistically significant results are indicated by asterisks (after Bonferroni corrections).

| Dependent<br>variables | Type of<br>lights | Signal-no<br>signal | Comparisons<br>between light<br>types | No<br>signal | Signal |
| --- | --- | --- | --- | --- | --- |
| Time spent in<br>movement (%) | Darkness | 0.340 | Darkness-HPS | 0.014* | 0.012* |
|  | HPS | 0.388 | HPS-LED | 1 | 1 |
|  | LED | 0.349 | LED-Darkness | 0.003* | 0.003* |
| Time spent in<br>shelter (%) | Darkness | 0.153 | Darkness-HPS | 1 | 0.185 |
|  | HPS | 0.008* | HPS-LED | 1 | <0.001* |
|  | LED | 0.139 | LED-Darkness | 1 | 0.006* |

**Figure 1.**
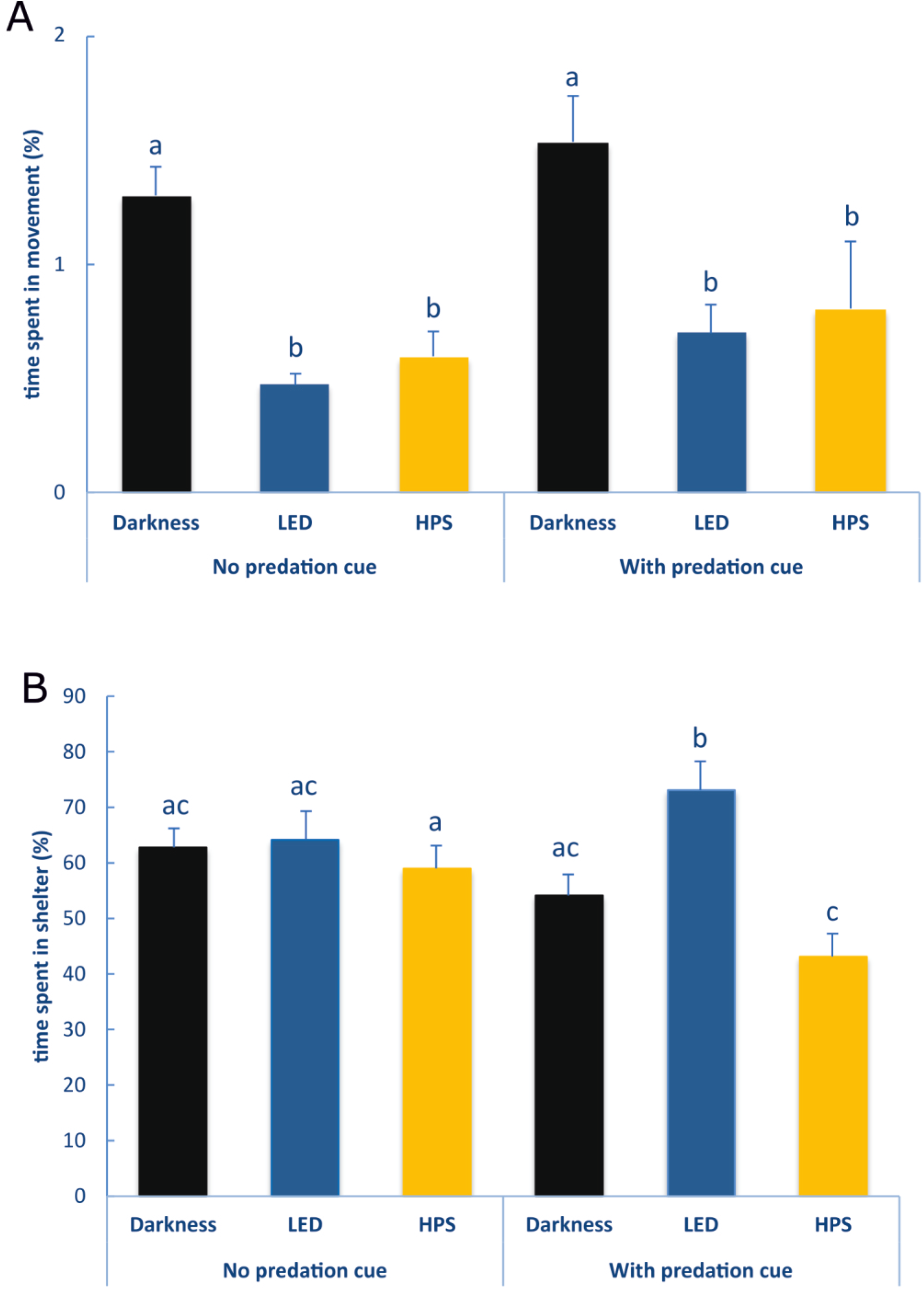
Effects of different types of light and predation cues on (A) time spent in movement and (B) time spent in shelter by caddisflies (tested with GLM). Bars labelled with different letters indicate statistically significant differences.

## Discussion

Our findings revealed that behavior of caddisflies differed between dark night-time conditions and artificially lit conditions. Moreover, their response to ALAN was modified by the presence of predation cues depending on the spectral composition of the light.

Caddisflies exposed to ALAN without predation cues reduced activity levels compared to darkness, regardless of the light type. This indicates that lower mobility was the result of overall increased light levels at night rather than specific spectral composition of ALAN. A similar behavioral response to nocturnal illumination was observed in nocturnal freshwater amphipods which reduced locomotor activity, but did not discriminate between warm LED and HPS light (Czarnecka et al. 2021). Studies conducted in marine environment also supported observations that nocturnal invertebrates often respond to night lighting by limiting their activity (Duarte et al. 2019; Manríquez et al. 2019). Since increased light levels at night could enhance the visibility of invertebrates, reducing movements may be an effective strategy impeding their detection by potential predators. Some studies also showed that invertebrates not only reduced their activity under nocturnal illumination, but also spent more time in shelters (Fischer et al. 2020; Czarnecka et al. 2021). However, in our experiment we found no relationship between ALAN exposure and shelter use in caddisflies. This may be due to the fact that *L. hirtum* builds cases consisted of sand grains, which are a form of physical protection, also against predation (Hansell 1972). Additionally, dark colours of their cases makes them less conspicuous in the wild as they are often difficult to distinguish them from the background.

The appearance of injured conspecifics in the immediate vicinity provides information about potential predation risk and usually induces anti-predatory responses such as reduced locomotor activity or hiding (Wisenden et al. 2001; Schoeppner and Relyea 2009; Smith and Webster 2015). Alarm cues also trigger other defensive strategies in animals, e.g. grouping or dispersal of conspecifics, providing benefits such as reduced capture risk and confusion of predators (Foster and Treherne 1981; Godin 1986; Morgan and Colgan 1987; Tosh et al. 2006; Roberts 1996). Therefore, we expected that predation cues would also affect the caddisfly behavior in dark-night conditions in our study. However, we found no impact of predation cues on both caddisfly activity and shelter use compared to the treatment without cues. Similarly, alarm cues did not influence caddisfly aggregation or dispersal. It is possible that the lack of response of caddisflies to predation cues in darkness may also be attributed to the presence of protective cases. The cases have been shown to increase the survival of caddisflies when encountering predators (Ferry et al. 2013), therefore this may explain their sense of safety despite the presence of predation cues.

The caddisfly response changed significantly when ALAN and predator cues occurred simultaneously, and this effect depended on the spectral quality of the light. The combination of exposure to LED light and predation cues increased defensive behaviors in caddisflies. They not only decreased their activity, but also spent more time in shelter compared to HPS light and darkness. LED light is characterized by a greater contribution of blue wavelengths than HPS light (Davies et al. 2013). Several studies have shown that some aquatic species are particularly sensitive to blue enriched LED light. Increased avoidance of cool LED light was found in river crayfish (*Cambarus chasmodactylus*) and spiny stream crayfish (*Faxonius cristavarius*) (Fischer et al. 2020), as well as in some amphipods (*Gammarus jazdzewskii*) (Czarnecka et al. 2022). Although information on the responses of caddisfly larvae to different types of light is limited, Nuntakwang et al. (2021) found that adults living in the terrestrial environment have the capability to perceive blue light. Another study demonstrated that caddisfly adults were particularly attracted by light traps equipped with metal-halide and LED lamps emitting light at short wavelengths (in the range of 360 to 407 nm) (Szanyi et al. 2022). Thus, terrestrial adult caddisflies and their immature aquatic stages responded differently to LED illumination, but in both cases this type of light disrupted their behavior.

Interestingly, caddisflies exposed to HPS light and predation cues spent less time in shelter than in the absence of the cues, although they showed similar, low levels of activity. This is unclear why the addition of predation cues triggered such behavior. Perhaps this may be a maladaptive response to predation threat under the influence of a novel anthropogenic factor. Nevertheless, such behavior may have a detrimental effect on the survival of caddisflies due to their increased visibility to potential predators (e.g. visually oriented fish) in ALAN (Czarnecka et al. 2019). Since hiding is usually more effective anti-predator strategy than merely limiting activity, caddisflies in HPS light may be more vulnerable to predation risk.

## Conclusions

Our research findings revealed that ALAN had a significant impact on the sheltering behavior and mobility of caddisflies. Moreover, we found that the presence of predation threat, particularly in combination with exposure to LED light, intensified the negative behavioral response of caddisflies. In such conditions, invertebrates not only decreased their activity but also spent more time in shelter compared to other treatments.

Our results highlight the importance of considering the combined effects of ALAN and biotic factors such as predation risk, because organisms are typically exposed to multiple stressors. The presence of predators, which is common in nature, can alter the response of caddisflies exposed to ALAN. Reduced activity of caddisfly larvae under predation risk and their tendency to stay in refuges in LED light may decrease their availability to potential predators, but on the other hand may also limit their involvement in other activities, e.g. searching for food, shredding leaf litter. Given the increasing use of LEDs in street lighting, also in suburban areas (van Langevelde et al. 2011; Pawson and Bader 2014; van Grunsven et al. 2014), a higher impact of ALAN on nocturnal invertebrates, including Trichoptera, should be expected.

## Notes

### Competing Interest Statement

The authors have declared no competing interest.

